# Polar relaxation by dynein-mediated removal of cortical myosin II

**DOI:** 10.1101/552372

**Authors:** Bernardo Chapa-y-Lazo, Motonari Hamanaka, Alexander Wray, Mohan K. Balasubramanian, Masanori Mishima

## Abstract

Nearly 6 decades ago, Lewis Wolpert proposed the relaxation of the polar cell cortex by the radial arrays of astral microtubules as a mechanism for cleavage furrow induction (White and Borisy, 1983; Wolpert, 1960). While this mechanism has remained controversial (Rappaport, 1996), recent work has provided evidence for polar relaxation by astral microtubules (Chen et al., 2008; Dechant and Glotzer, 2003; Foe and Dassow, 2008; Murthy and Wadsworth, 2008; Werner et al., 2007), although its molecular mechanisms remain elusive. Here, using *C. elegans* embryos, we show that polar relaxation is achieved through dynein-mediated removal of myosin II from the polar cortexes. Mutants that position centrosomes closer to the polar cortex accelerated furrow induction whereas suppression of dynein activity delayed furrowing. We provide evidence that dynein-mediated removal of myosin II from the polar cortexes triggers cortical flow towards the cell equator, which induces the assembly of the actomyosin contractile ring. These studies for the first time provide a molecular basis for the aster-dependent polar relaxation, which works in parallel with equatorial stimulation to promote robust cytokinesis.

## Results and Discussion

During animal cell cytokinesis, cleavage by constriction of an actomyosin contractile ring is spatially coupled to chromosome segregation by the mitotic apparatus (D’Avino et al., 2015; Green et al., 2012). The best-understood mechanism for spatial coupling is chemical signaling via activation of Rho GTPase at the cell equator by the central spindle, which is mediated by centralspindlin, a microtubule-bundling signaling hub, and ECT2 RhoGEF under the regulation of mitotic kinases (Mishima, 2016; White and Glotzer, 2012). A phosphatase-mediated negative signal from the kinetochores has also been reported to reduce the F-actin levels at the polar cortexes (Rodrigues et al., 2015). However, in cells with a relatively small mitotic spindle such as the *C. elegans* one-cell stage embryo, neither positive nor negative signals from the spindle or chromosomes alone can effectively work on the distant cell cortex at the initial stage of cytokinesis before the cleavage furrow deepens (Dassow, 2009; Mishima, 2016). It is known that astral microtubules play an important role in cleavage furrow induction (Baruni et al., 2008; Bringmann and Hyman, 2005; Bringmann et al., 2007; Dechant and Glotzer, 2003; Werner et al., 2007), but the molecular details have remained unclear.

Nonmuscle myosin II is a major component of the cortical actomyosin network and the cytokinetic contractile ring, and is crucial for animal cell cytokinesis. Myosin dynamics in dividing *C. elegans* embryos has previously been studied, but data are limited to a low temporal resolution (>∼10 s) for a 3D volume (Carvalho et al., 2009; Jordan et al., 2016; Khaliullin et al., 2018) or, at higher temporal resolution, to the cell surface (Reymann et al., 2016; Werner et al., 2007). To examine the rapid dynamics of myosin II on the mitotic spindle and astral microtubules, we performed fast time-lapse recording (∼every second) of GFP-tagged nonmuscle myosin II, expressed from the endogenous locus (*nmy-2(cp13)*) (Dickinson et al., 2013), imaging a 2 µm thick z-section at the embryo midplane with 0.5 µm z-steps. Embryos within the eggshell were immobilized on the surface of a cover glass without any deformation. During metaphase, in addition to accumulation at the cell cortex, NMY-2::GFP was detected on the spindle (**Fig. 1a** metaphase, **Supplementary Video 1**). Although cytoplasmic particulate signals were also detected, they only showed diffusive random motion and did not show directional movement. By contrast, after anaphase onset, unidirectional movement of NMY-2 particles towards the spindle poles was detected (**Fig. 1a** anaphase, **Supplementary Video 2**), with a mean velocity of ∼0.7 µm/s (**Fig. 1b, c**). In instances where we could clearly see internalization of NMY-2, we noticed that flanking NMY-2 signals moved apart (**Fig. 1d**), reminiscent of cortical relaxation after laser ablation (Mayer et al., 2010). This observation suggested that removal of NMY-2 from the cortex leads to a local reduction of the cortical tension/contractility.

**Figure 1.**
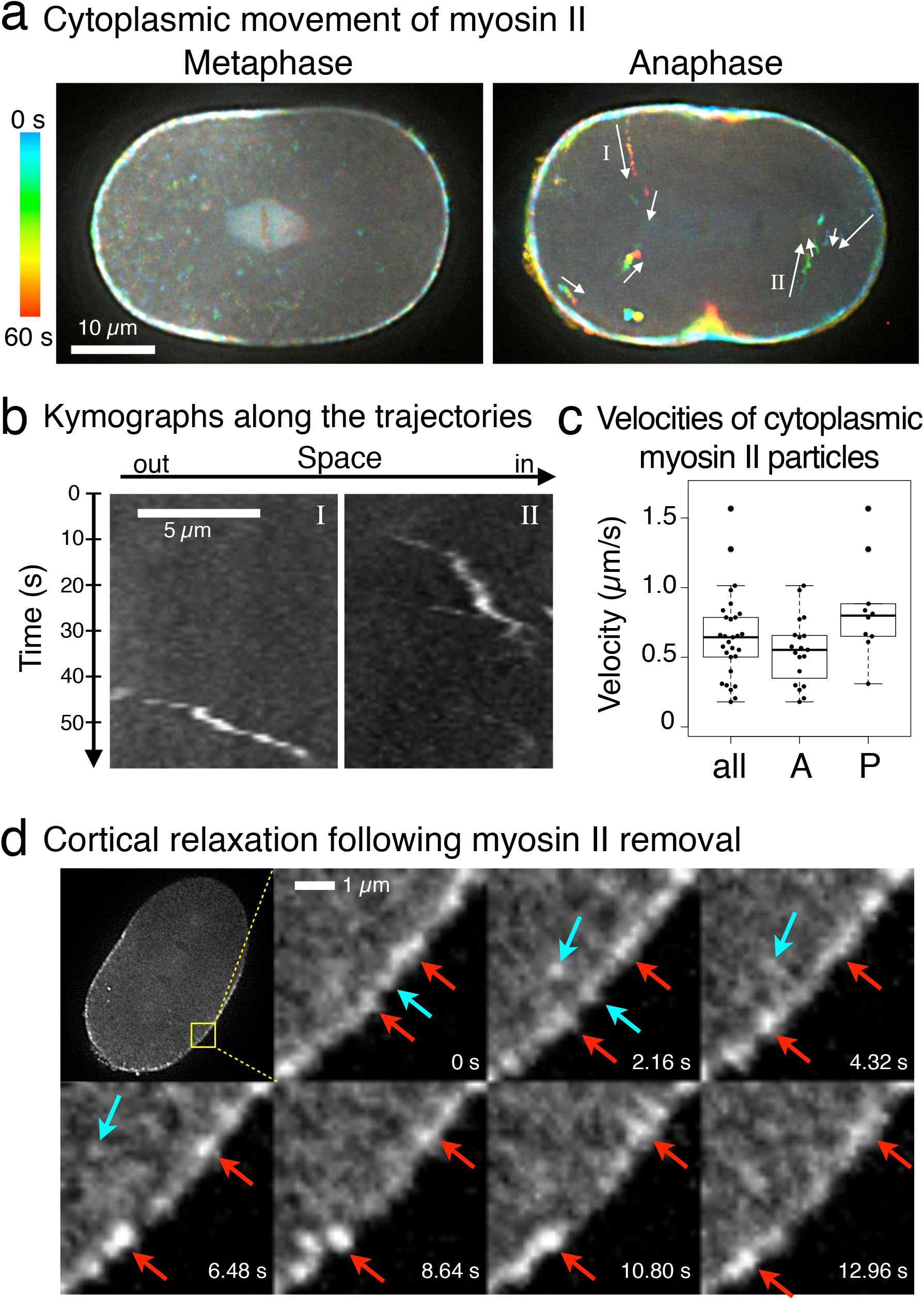
Removal of myosin II by centrosome-directed movement relaxes the cell cortex. **a,** GFP-tagged non-muscle myosin II (NMY-2) in the mid-plane of *C. elegans* embryos was imaged by spinning disk confocal microscopy every 0.83 s and is presented with temporal color-coding. **b,** Kymographs of the myosin II particles labeled I and II in the anaphase embryo in panel **a**. **c,** Velocity distributions of the unidirectional movement of the cytoplasmic myosin II particles found in anterior (A) and posterior (P) sides of anaphase embryos (n=19 and 10, respectively, from 4 embryos). **d,** Time lapse of a section of the anterior cortex at a higher magnification. Cortical patches of NMY-2 (blue arrows) were sequentially internalized at around 1 s and 3 s. Following this, the distance between the flanking patches/particles (red arrows) gradually increased, indicating the local relaxation of the cortical actomyosin network.

The direction and velocity of cytoplasmic movement of NMY-2 were consistent with motility by dynein, a microtubule minus-end-directed motor (Reck-Peterson et al., 2018). To test the role of microtubules in the movement of NMY-2, embryos were treated with a microtubule depolymerizing drug, nocodazole. As expected, the cytoplasmic movement of NMY-2 particles was quickly suppressed after nocodazole treatment, while it was not affected at all in control embryos treated with the drug vehicle (**Fig. 2a**). To test the role of dynein in the directional transport of myosin II as its cargo, we used RNAi to deplete embryos of the dynein heavy chain DHC-1 (Gönczy et al., 1999) or to deplete LIN-5, the ortholog of a dynein regulator NuMA (Lorson et al., 2000; Radulescu and Cleveland, 2010). We also tested ciliobrevin D, an inhibitor of vertebrate cytoplasmic dynein, but this did not cause any mitotic or developmental abnormalities. No unidirectional cytoplasmic movement of NMY-2 was observed in the *dhc-1(RNAi)* embryos (**Fig. 2b, Supplementary Video 3**), which underwent massively disorganized cell division reflecting the multiple mitotic functions of dynein (functions in centrosome separation, metaphase chromosome congression, spindle positioning and elongation, and chromosome segregation) (Gönczy et al., 1999). In the *lin-5(RNAi)* embryos, mitosis proceeded more normally except for expected defects in spindle positioning and elongation (Galli et al., 2011; Lee et al., 2015; Panbianco et al., 2008; Srinivasan et al., 2003) but still no cytoplasmic unidirectional movement of NMY-2 particles was detected (**Fig. 2c, Supplementary Video 4**). While unidirectionally moving NMY-2 particles in the cytoplasm gradually increased during anaphase in control embryos, such particles were essentially undetectable in the embryos depleted of LIN-5 or its binding partners, GPR-1/2, the Pins/LGN orthologs (**Fig. 2d**). These results indicate that the movement of NMY-2 from the cortex towards the spindle poles depends on astral microtubules and is driven by dynein.

**Figure 2.**
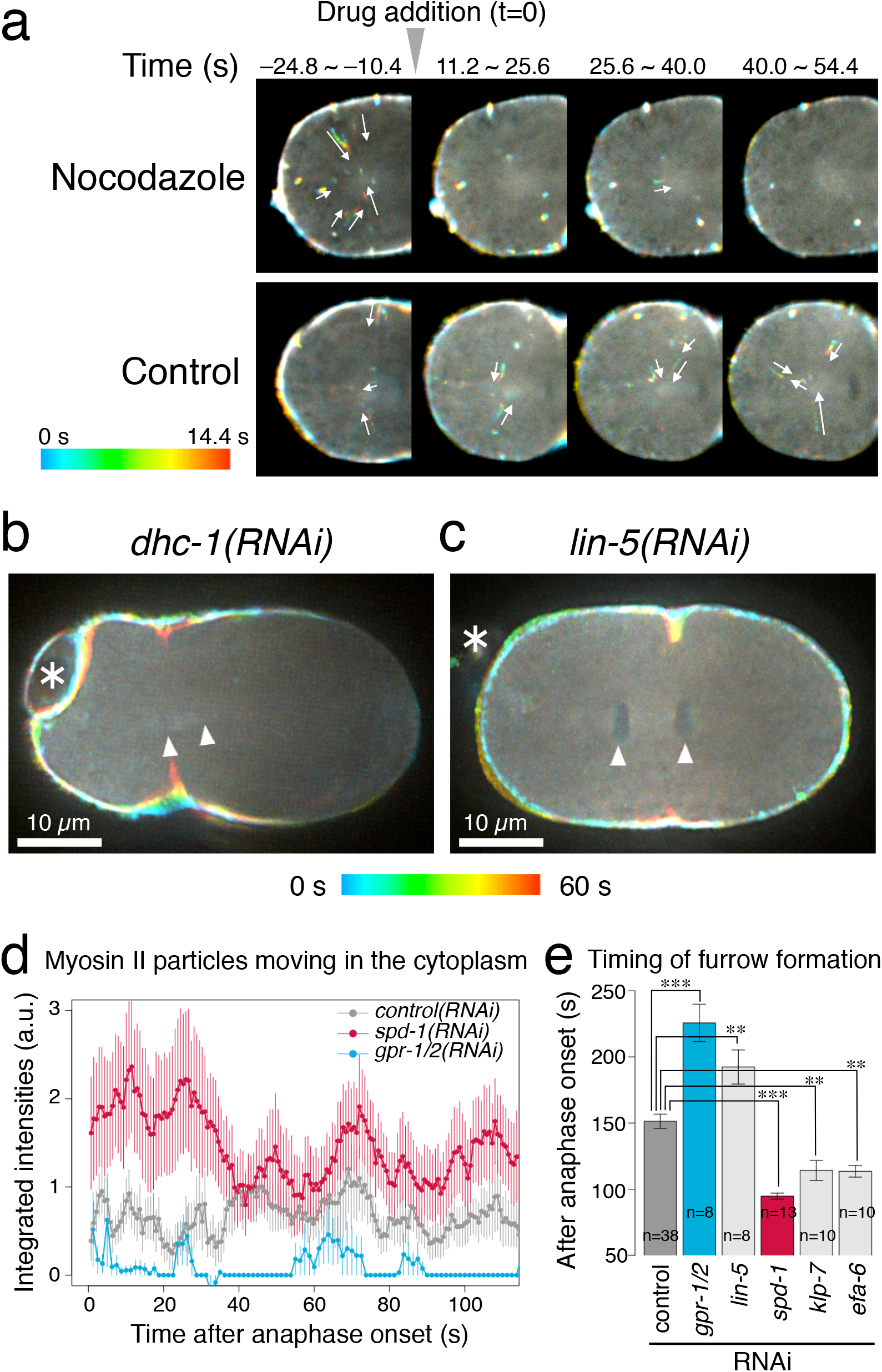
Astral microtubules and dynein drive internalization of myosin II. **a,** Embryos expressing NMY-2::GFP were treated with 15 µM nocodazole, a microtubule poison, or the drug vehicle (control) during anaphase (at 0 s). Nocodazole strongly suppressed the cytoplasmic motility of myosin II. **b, c,** Depletion of the cytoplasmic dynein heavy chain DHC-1 (**b**) or the dynein regulator NuMA (LIN-5) (**c**) eliminated the cytoplasmic unidirectional movement of myosin II. **d,** Quantification of the myosin II particles that show unidirectional motility in the cytoplasm of the control, *spd-1(RNAi)* and *gpr-1/2(RNAi)* embryos (n=22, 23 and 12, respectively). The average of the sum of the particles’ intensities across embryos for each time point was plotted with the standard error. Depletion of the LIN-5 partner GPR-1/2 abolished the cytoplasmic movement of myosin II particles, while depletion of SPD-1/PRC1, which accelerates the approach of the asters to the cortex (**Fig. 3a, b**), promoted an earlier internalization of myosin II (refer to the **Fig. 3c** as well). **e,** Timing of the initial sign of furrow formation determined by visual inspection of anonymized movies (mean ± s.e.). Statistical significance was tested by linear modeling with correction for multiple comparisons by Dunnett’s method. ** and *** indicate p<0.01 and <0.001, respectively.

In various cell types, including *C. elegans* embryos (Dechant and Glotzer, 2003; Khaliullin et al., 2018; Werner et al., 2007), echinoderm embryos (Dassow et al., 2009; Foe and Dassow, 2008), silkworm spermatocytes (Chen et al., 2008), and mammalian cultured cells (Murthy and Wadsworth, 2008), evidence has been accumulating for the suppression of the contractility of the polar cortexes by radial arrays of astral microtubules (polar relaxation). The polar relaxation contributes to furrow formation often by promoting the flow of the actomyosin network within the cell cortex (cortical flow) (Chen et al., 2008; Dan, 1954; Khaliullin et al., 2018; Murthy and Wadsworth, 2008; Werner et al., 2007), which contributes to the assembly of the actomyosin contractile ring (DeBiasio et al., 1996; Reymann et al., 2016; Salbreux et al., 2009; Turlier et al., 2014; Zhou and Wang, 2008), and also by releasing cytoplasmic hydrostatic pressure, which otherwise destabilizes the furrow (Sedzinski et al., 2011). The local reduction of the cortical contractility upon removal of NMY-2 from the cortex (**Fig. 1d**) caused us to hypothesize that the dynein-driven transport of myosin II along astral microtubules might be the major mechanism for this aster-dependent polar relaxation. If so, suppression of the dynein-mediated internalization of NMY-2 should interfere with furrow formation. Indeed, we observed a significant delay in the furrow formation in the embryos depleted of LIN-5 or GPR-1/2 and thus lacking the NMY-2 internalization (**Fig. 2e**). While the first sign of furrow ingression was detected by visual inspection at ∼150 s post anaphase onset (p.a.o.) in the control embryos, it was delayed by more than 40 s in the *lin-5(RNAi)* and *gpr-1/2(RNAi)* embryos (**Fig. 2e**). These data suggest a substantive role for dynein-mediated myosin II transport in cleavage furrow formation.

To further test the role of the dynein-mediated NMY-2 internalization in the cleavage furrow formation, we sought to enhance it. It has been shown that the density of astral microtubules decreases as the distance from a pole increases (Dechant and Glotzer, 2003). This means that bringing the centrosomes closer to the cell cortex should upregulate the dynein-mediated removal of the cortical myosin II by astral microtubules. This is achievable, for example, by disrupting the central spindle, which mechanically links the two spindle poles and thus opposes spindle pole separation. The central spindle can be broken by depletion of SPD-1 (Verbrugghe and White, 2004), the ortholog of vertebrate PRC1, a highly conserved microtubule-bundling protein crucial for the central spindle formation (Jiang et al., 1998; Mollinari et al., 2002; Subramanian et al., 2013).

In normal embryos, the spindle elongates from ∼12 µm to ∼24 µm in ∼200 s after anaphase onset mainly by the action of cortical pulling forces acting on the spindle poles, in coordination with microtubule-sliding and polymerization at the spindle midzone (Grill et al., 2001; Lee et al., 2015; Maton et al., 2015; Nahaboo et al., 2015; Panbianco et al., 2008; Pecreaux et al., 2006; Saunders et al., 2007). Due to the asymmetry of the pulling forces applied to the spindle, the posterior pole shows faster posterior-directed translational movement and a larger oscillatory movement perpendicular to this, in comparison to the anterior pole. It reaches within 9 µm from the posterior polar cortex in 60 s p.a.o (**Fig. 3a, b** *control(RNAi)*, and **Supplementary Fig. 1** *control(RNAi)* for more time points and cortical position vs distance plots). This creates a local minimum in the pole-to-cortex distance, where the influence of the radial array of astral microtubules locally maximizes, at the posterior tip of the embryo. On the other hand, the anterior pole largely stays around its initial position for ∼70 s before it starts to move towards the anterior. As a consequence, the pole-to-cortex distance of the anterior cortex remained at similar levels to or higher than that of the equatorial cortex until ∼95 s p.a.o. although a zone of a shallow minimum is formed around 10 µm anterior to the future cleavage site (**Fig. 3a, b** and **Supplementary Fig. 1** *control(RNAi)*). Importantly, the movements of the two spindle poles towards the opposite polar cortexes created a local maximum in the pole-to-cortex distance at the equatorial region (∼3 µm posterior from the exact center of the A-P axis), which predicts the site of cleavage furrow ingression.

**Figure 3.**
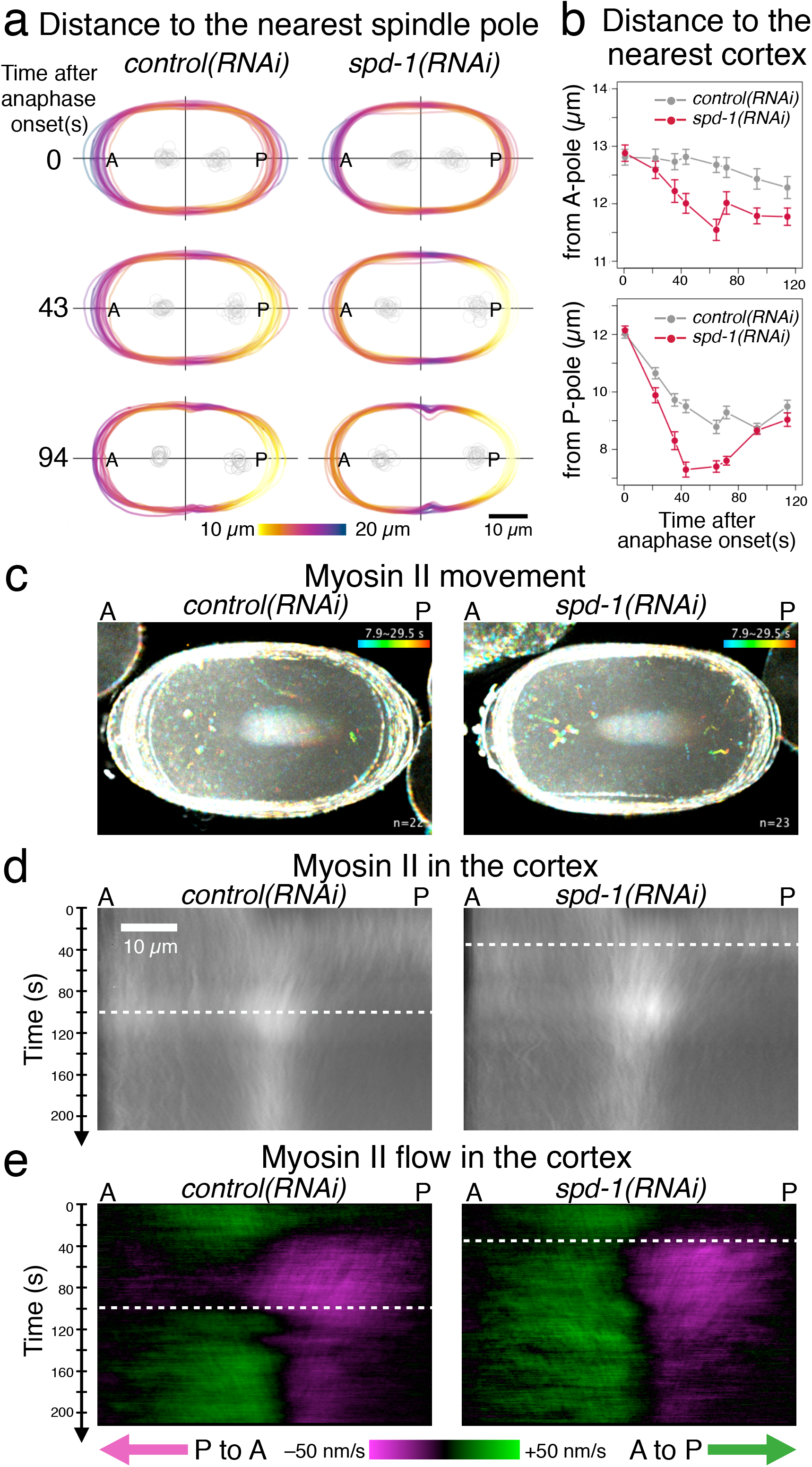
The spindle pole-cortex geometry regulates cleavage furrow formation by controlling the distribution and flow of cortical myosin II. **a and b,** Distance between the spindle poles and the cell cortex. In **a**, the distance from each point on the cell periphery to the nearest spindle pole was measured and presented with color coding. The positions of the spindle poles are illustrated in gray. In **b**, the distance from the anterior pole (A-pole) or the posterior pole (P-pole) to the nearest cortex was measured. Rupture of the central spindle by depletion of the key microtubule-bundler SPD-1/PRC1 accelerated the spindle’s pole-to-pole elongation, and thus, the decrease in pole-to-cortex distance. **c,** Effect of depletion of SPD-1 on the cytoplasmic myosin II particles. Temporal color-coded trajectories of NMY-2::GFP from control (n=22) or *spd-1(RNAi)* (n=23) embryos were overlaid by maximum projection. Moving myosin II particles appeared earlier in the *spd-1(RNAi)* embryos. Refer to **Fig. 2d** as well. **d and e,** Density (**d**) and flow (**e**) of myosin II in the cell cortex. **d,** Cortical NMY-2::GFP was quantified along the cell periphery from the anterior tip to the posterior tip and normalized with the local background and cytoplasmic levels. Average across the embryos from two sets of recordings (22 control and 23 *spd-1(RNAi)* embryos for 0 to 140 s p.a.o. and 56 control and 19 *spd-1(RNAi)* embryos for 70 to 210 s p.a.o.) were merged and presented as a kymograph for each condition (see Materials and Methods for more details). **e,** The flow of NMY-2::GFP along the cell periphery was computed by one-dimensional particle image velocimetry and is presented as a kymograph. The green signal indicates flow towards the posterior and the magenta towards the anterior. The timing of the transition of the cortical flow into the bidirectional mode (white dashed line) is accelerated in the embryos depleted of SPD-1, in which the spindle asters are located closer to the cortex.

In *spd-1(RNAi)* embryos, the anterior and posterior asters are separated from each other immediately after anaphase onset as a consequence of mitotic spindle rupture (Lee et al., 2015; Verbrugghe and White, 2004). This accelerated the approaching of both the poles to the cortex, positioning both of them closer to the cortex than the final levels in the control embryos within 40 s after anaphase onset (**Fig. 3a, b** and **Supplementary Fig. 1** *spd-1(RNAi)*). This established a deeper pole-to-cortex distance minimum in the anterior cortex earlier than in control embryos (**Fig. 3 a, b** and **Supplementary Fig. 1** *spd-1(RNAi)*). As expected, the cytoplasmic myosin particles moving unidirectionally along microtubules appeared earlier in the *spd-1(RNAi)* embryos than in controls (**Figs. 2d** and **3c**). Importantly, despite the disruption of the central spindle, an important source of a positive signal for the contractile ring assembly, the initiation of cleavage furrow formation was accelerated by 50 s in the *spd-1(RNAi)* embryos (**Fig. 2e**). A similar but slightly milder acceleration of furrow initiation was observed by milder acceleration of the pole-to-pole separation (spindle elongation) by depletion of KLP-7, a microtubule depolymerizer that targets astral microtubules (Grill et al., 2001; Han et al., 2015; Rankin and Wordeman, 2010; Srayko et al., 2005), or by depletion of EFA-6, a negative regulator of the dynein-dependent aster-cell cortex interaction (O’Rourke et al., 2010) (**Fig. 2e**). These observations further support the role of dynein-mediated myosin II internalization by astral microtubules in the induction of the cleavage furrow.

Next, we investigated how dynein-mediated internalization of myosin II controls cleavage furrow formation. In order to track cortical myosin II dynamics in relation to the geometry of the spindle, we measured the NMY-2::GFP signal along the cell periphery from the anterior tip to the posterior tip at each time point (for details, see **Materials and Methods** and **Supplementary Fig. 2**). The average across embryos was displayed as a kymograph representing the dynamics of the distribution of the cortical myosin II (**Fig. 3d**). Using the same data, the cortical flow along the cell periphery was computed in individual embryos by one-dimensional particle image velocimetry (PIV), which gave comparable values to the anterior-posterior component of the flow velocity obtained by two-dimensional PIV (Khaliullin et al., 2018; Reymann et al., 2016; Sugioka and Bowerman, 2018). The average across embryos was displayed as a kymograph in which the green signal indicates flow from the anterior to the posterior, and the magenta indicates flow from the posterior to the anterior (**Fig. 3e**).

In normal embryos, during metaphase, NMY-2 showed an asymmetric cortical localization slightly enriched at the anterior cortex (**Fig. 3d** control, 0 s), as a remnant of post-fertilization polarization (Munro et al., 2004; Werner et al., 2007). After anaphase onset, the signal in the posterior cortex gradually increased and subsequently the global signal reached a uniform distribution at around 20 s p.a.o. (**Fig. 3d** control, 0∼20 s). This state was maintained until 40 s p.a.o., when the posterior pole approached within ∼10 µm from the posterior cortex, forming a minimum at the posterior pole. At this point, the signal in the posterior cortex started to decline (**Fig. 3d** control, ∼40 s) and, simultaneously, a flow from the posterior to the anterior appeared, which gradually became stronger and extended towards the anterior cortex until ∼100 s p.a.o. (**Fig. 3e** control, 40∼100 s, the magenta signal on the right half of the panel). This resulted in a broad and moderate accumulation of NMY-2 at the equatorial zone (**Fig. 3d** control, ∼100 s p.a.o.). By contrast, the signal on the anterior cortex stayed largely constant and without a strong flow (**Fig. 3d and e**, control, 40∼100 s, the left half of the panels). Meanwhile, the anterior spindle pole started to move slowly towards the anterior tip of the embryo. Following this period, around 100 s p.a.o., a drop in the NMY-2 signal was observed in the anterior cortex (**Fig. 3d**), concomitantly with the appearance of a strong flow towards the posterior (**Fig. 3e** control, >100 s, green signal on the left half of the panel). This resulted in a period of bidirectional flows, in which the flow towards the anterior from the posterior cortex converged with the flow towards the posterior from the anterior cortex at the cell equator. The bidirectional flows gradually sharpened the equatorial zone of NMY-2 accumulation (**Fig. 3d** control) and resulted in furrow formation around 150 s. p.a.o (**Fig. 2e**).

As described above, depletion of SPD-1 accelerates the approach of the centrosomes to the polar cortexes and promotes the dynein-mediated internalization of myosin II. This caused a drastic change in the dynamic behavior of NMY-2 at the cortex. Following the earlier appearance of the anterior minimum in the pole to cortex distance (**Fig. 3b** *spd-1(RNAi)* 20 and 40 s), a drop of NMY-2 signal at the anterior cortex was observed at 60 s p.a.o. (**Fig. 3d** *spd-1(RNAi)*, 60 s p.a.o.), accompanied by an earlier rise of a stronger posterior-directed flow in the anterior cortex (**Fig. 3e** *spd-1(RNAi),* 60 s p.a.o.). This accelerated entry into the phase of bidirectional flows by ∼50 s (**Fig. 3e**, dashed white lines) and resulted in earlier and stronger accumulation of myosin II at the equator (**Fig. 3d**) together with furrow formation ∼60 s earlier than in control embryos (**Fig. 2e**). Consistent with this, in the *efa-6(RNAi)* and *klp-7(RNAi)* embryos, in which the approaching of the spindle poles to the polar cortexes is mildly accelerated, the bidirectional flow (**Supplementary Fig. 3**) and the furrow ingression (**Fig. 2e**) occurred 20∼30 s earlier than in control embryos. These observations indicate that removal of myosin II from a polar cortex by dynein and astral microtubules triggers a cortical flow away from it, and that creation of bidirectional flows towards the equator plays a key role in the cleavage furrow formation.

Depletion of the dynein regulators LIN-5 and GPR-1/2 completely eliminates the dynein-driven internalization of myosin II from the cell cortex (**Fig. 2c and d**). Since these factors also have a role in spindle positioning by the cortical pulling forces (Colombo et al., 2003; Gotta et al., 2003; Pecreaux et al., 2006; Srinivasan et al., 2003), in the *lin-5(RNAi)* or *gpr-1/2(RNAi)* embryos, at anaphase onset, the mitotic spindle is positioned slightly more anterior than in control embryos, and after anaphase onset, the posterior shift, spindle elongation, and spindle oscillation are all suppressed. Although depletion of these dynein regulators does not itself prevent furrow formation, combinations with defects in central spindle factors such as SPD-1/PRC1 or ZEN-4/MKLP1 cause synthetic failure of furrow formation (Bringmann et al., 2007; Dechant and Glotzer, 2003; Maton et al., 2015; Verbrugghe and White, 2007; Werner et al., 2007). These observations indicate that dynein and these dynein regulators work in furrow induction in parallel with the central spindle/centralspindlin-dependent pathway. However, it has remained unclear whether dynein and its regulators positively control cortical contractility by equatorial stimulation or negatively by polar relaxation. Considering the novel mechanism of centrosome-directed transport of myosin II by dynein that we have discovered, it is highly likely that these factors work in polar relaxation and not in equatorial stimulation. To test this and to clarify the contributions of the dynein-dependent and centralspindlin-dependent pathways, we assessed the effect of depletion of GPR-1/2 on the dynamics of cortical myosin II and examined the synthetic effect with the inhibition of the centralspindlin-dependent equatorial stimulation observed in the *cyk-4(or749)* mutant embryos (**Fig. 4a-c**). The point mutation of this temperature sensitive allele lies in the GAP domain of CYK-4 (Canman et al., 2008; Davies et al., 2014) and prevents the equatorial stimulation since the GAP domain plays a role in equatorial stimulation by inactivation of Rac (Canman et al., 2008; Zhuravlev et al., 2017) or by activation of Rho via the RhoGEF ECT-2 (Loria et al., 2012; Tse et al., 2012; Zhang and Glotzer, 2015). Importantly, unlike other methods used to perturb the centralspindlin/central spindle-dependent pathway, this allele, at the restrictive temperature (25 °C), achieves this without affecting the formation and maintenance of the central spindle or the localization of centralspindlin to the spindle midzone (Canman et al., 2008).

**Figure 4.**
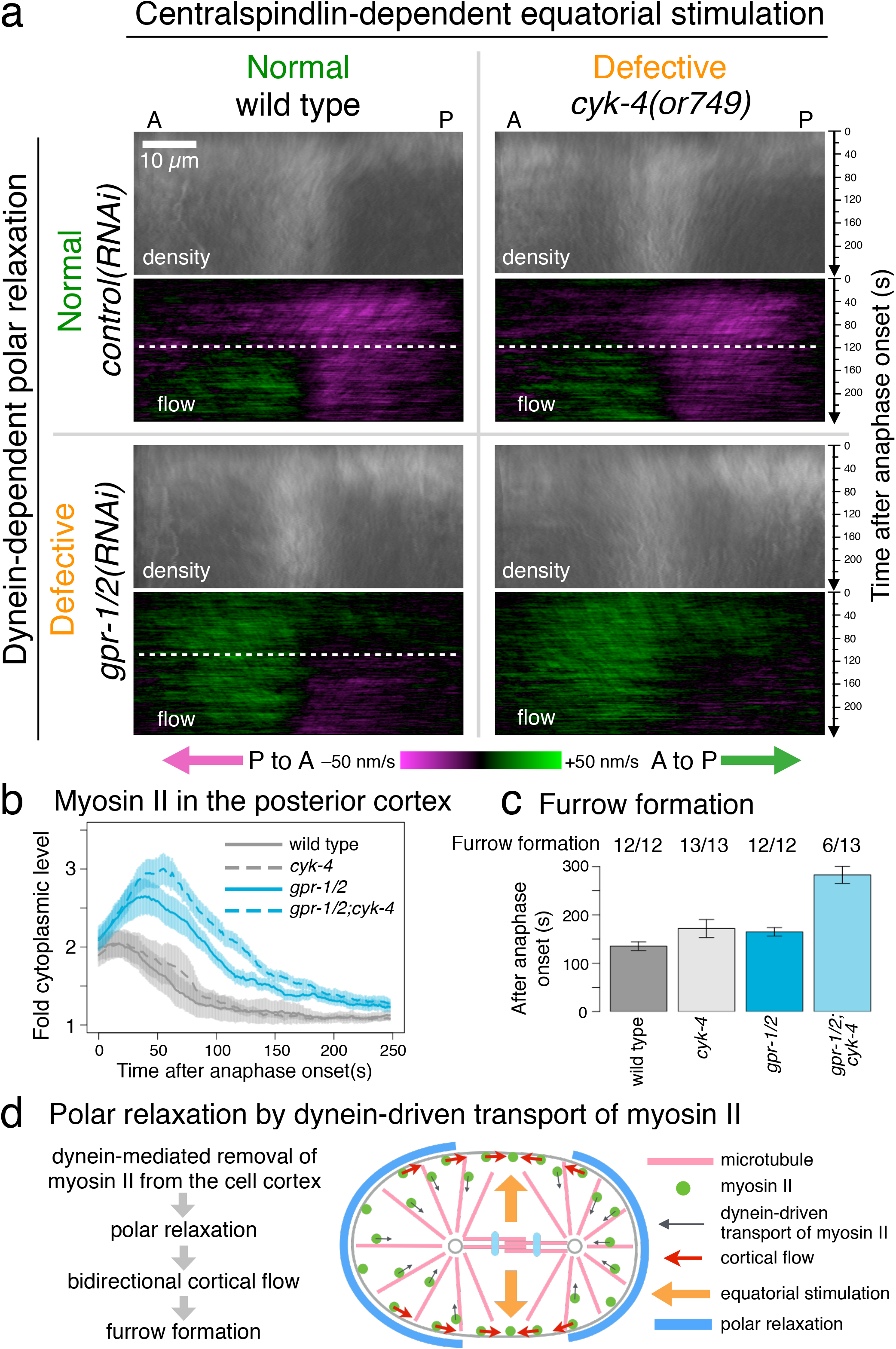
Polar relaxation by dynein-mediated removal of cortical myosin II induces furrow formation in parallel with equatorial stimulation. **a∼c,** Synthetic effect of the simultaneous inhibition of dynein activity (*gpr-1/2(RNAi)*) and centralspindlin-mediated equatorial stimulation (*cyk-4(or749)*) on the distribution and cortical flow of NMY-2::GFP (**a, b**) and the timing of furrow formation (**c**). Since the *cyk-4(or749)* mutation is temperature-sensitive, embryos were imaged at 25°C. **b,** Temporal change of the density of myosin II in the posterior cortex within 10 µm from the posterior tip was plotted (mean across embryos ± s.e.). The defect in the centralspindlin-dependent equatorial stimulation (*control(RNAi);cyk-4(or749),* top right in **a.**) did not severely affect the patterns of the myosin II distribution and the cortical flow, which relied on the dynein-dependent removal of myosin II (*gpr-1/2(RNAi);wild type,* bottom left in **a.**). Inhibition of both the centralspindlin-dependent pathway and the dynein-dependent pathway almost completely abolished the bidirectional flows (**a, b**) and severely affected the furrow formation (**c**). **d,** Schematic showing how bidirectional cortical flow triggered by dynein-dependent removal of myosin II from the polar cortexes leads to cleavage furrow formation. This is crucial for the initial induction of the cleavage furrow before the equatorial stimulation from the central spindle becomes effective.

In the embryos depleted of GPR-1/2 in the wild type background, NMY-2::GFP showed hyperaccumulation to the posterior cortex during early anaphase (**Fig. 4a**, *gpr-1/2(RNAi)*;wild type, “density”, and **Fig. 4b** *gpr-1/2*), confirming the role of dynein-driven transport of myosin II in the removal of myosin II from the cell cortex (polar relaxation). This was accompanied by a reversal of the global flow that lasts until 120 s p.a.o. (**Fig. 4a**, *gpr-1/2(RNAi)*;wild type, “flow”) and by delayed furrow ingression (**Fig. 4c** *gpr-1/2*). Although bidirectional flows appeared after 120 s p.a.o., the posterior-to-anterior flow towards the equator in the posterior cortex was much slower than in the wild type embryos. By contrast, in the *cyk-4(or749)* embryos, no obvious difference in the cortical distribution and the cortical flow of myosin II was observed during early anaphase (**Fig. 4a** *control(RNAi);cyk-4(or749)*). The transition into the bidirectional flow also occurred at similar timing to that in the wild type embryos (**Fig. 4a** dashed white lines), indicating a limited role of the centralspindlin-dependent equatorial stimulation in the regulation of the cortical flow during early anaphase. Interestingly, in the embryos defective for both the centralspindlin-dependent and dynein-dependent pathways, bidirectional flow was almost completely abolished (**Fig. 4a** *gpr-1/2(RNAi);cyk-4(or749)*). The *cyk-4* mutation further enhanced the delay of the clearance of myosin II from the posterior cortex due to the GPR-1/2 depletion (**Fig. 4a, b**). These results indicate that the weak bidirectional cortical flow observed in the *gpr-1/2(RNAi)* embryos in the wild-type background was caused by the centralspindlin-dependent equatorial stimulation. While all the embryos defective only for the centralspindlin-dependent pathway or the dynein-dependent pathway could initiate furrowing, albeit with some delays, about half of the embryos defective for both the pathways failed to form a furrow (**Fig. 4c**). The rest of the embryos could only form a very shallow furrow, which eventually regressed, after a long delay (**Fig. 4c**). Depletion of LIN-5 also prevented furrow formation or deepening in the *cyk-4* mutant background (**Supplementary Fig. 4**). Taken together, these data indicate that, in the *C. elegans* one-cell stage embryo, the polar relaxation triggered by dynein-driven myosin II transport contributes to the furrow induction in parallel with the centralspindlin-dependent equatorial stimulation (**Fig. 4d**), especially during the early establishment of the bidirectional cortical flows (**Figs. 3e and 4a**).

Since the first proposal of the polar relaxation by astral microtubules (Wolpert, 1960), its molecular mechanism has remained unclear. Here we revealed a fundamental mechanism in which centrosome-directed transport of myosin II along astral microtubules, driven by dynein, reduces the contractility of the polar cortex by removing myosin II from it (**Fig. 4d**). The local reduction of cortical contractility triggers a global cortical flow, and the geometry of the mitotic apparatus plays a key role in switching the flow from a unidirectional mode to a bidirectional mode. The bidirectional cortical flow towards the cell equator contributes to the equatorial accumulation of the actomyosin network and to the cleavage furrow formation (DeBiasio et al., 1996; Reymann et al., 2016; Salbreux et al., 2009; Turlier et al., 2014; Werner et al., 2007; Zhou and Wang, 2008). In addition to the central spindle, the molecules for equatorial simulation can be recruited to the anti-parallel overlaps of the equatorial astral microtubules (Argiros et al., 2012; D’Avino et al., 2006; Nguyen et al., 2014; Nishimura and Yonemura, 2006; Su et al., 2014; Uehara et al., 2016) and to the microtubule ends that laterally associate with the furrow cortex in monopolar cytokinesis (Canman et al., 2003; Hu et al., 2008; Kitagawa et al., 2013; Shrestha et al., 2012). It has also been reported that precise microtubule polymerization/depolymerization dynamics does not play a crucial role for furrow induction (Strickland et al., 2005). Considering these observations, we can summarize the action of microtubules on the nearby cell cortex into a simple rule: microtubules laterally associated with the cortex, which are often in an anti-parallel configuration, promote contractility, whereas microtubules pointing towards the cortex relax contractility.

How myosin II is linked to dynein is currently unclear. Whether and how regulators of the cortical pulling forces (Fielmich et al., 2018; Kiyomitsu and Cheeseman, 2013; Kotak et al., 2014; Rodriguez-Garcia et al., 2018; Schmidt et al., 2017; Sugioka and Bowerman, 2018) and cortical asymmetry (Chartier et al., 2011; Pacquelet et al., 2015; Sugioka et al., 2018), as well as membrane additions (Gudejko et al., 2012) or invaginations/endocytosis (Redemann et al., 2010; Tse et al., 2011), are involved in dynein-driven transport of myosin II (and, vice versa) will be important future questions. Compared to a mechanism that depends on a diffusive signal, direct remodeling of the actomyosin network through a physical interaction with microtubules has the advantage of keeping the effect precisely localized at a long distance. We speculate that the dynein-driven myosin II transport mechanism revealed in this work may have a broader biological role in controlling cell shape in other cell/tissue contexts, such as in cell migration and epithelial morphogenesis, since both dynein and myosin II are universally expressed in metazoan cells.

## Materials and Methods

### *C. elegans* strains and culture

The *C. elegans* strains used in this study are listed in Supplementary Table 1 and were maintained at 20°C, except for those containing the temperature-sensitive *cyk-4(or749)* allele, which were maintained at 15°C. To create the QM160 strain (*nmy-2(cp13[nmy-2::gfp + LoxP]) I; ltIs37[(pAA64) pie-1p::mCherry::his-58 + unc-119(+)] IV*), LP162 (*nmy-2(cp13)*) (Dickinson et al., 2013) and OD56 (*ltIs37*) (Essex et al., 2009) strains were crossed and double homozygous animals for both the *nmy-2(cp13)* and *ltIs37* alleles were isolated. The QM196 strain (*nmy-2(cp13[nmy-2::gfp + LoxP]) I; cyk-4(or749) III*) was generated by crossing LP162 and EU1404 (*cyk-4(or749)*) (Canman et al., 2008) strains. The *gpr-1/2* RNAi clone was from the Vidal Library (Rual et al., 2004) (Source BioScience). The *spd-1* RNAi clone was previously described (Lee et al., 2015). The other RNAi clones were from the Ahringer Library (Kamath et al., 2003) (Source BioScience). For RNAi in the non-temperature sensitive strains, L4 worms were plated on the RNAi plates and incubated at 20 °C for 48 h before dissection to obtain embryos. For RNAi in the temperature sensitive strains, L4 worms were plated and incubated at 16°C for 72 h.

### Live microscopy

For fluorescence live imaging, *C. elegans* embryos were immobilized on the cover glass surface of a 35 mm glass-bottom dish (Fluorodish, FD35, World Precision) and cultured in a drop of an osmolality controlled medium. Before experiments, the center of the cover glass was coated with 20 µl of 0.1 mg/ml poly-L-lysine (MW>300,000, Sigma P1524) for 1 h. After a quick wash with water, a circle was drawn with a PAP pen (Sigma, Z377821) around the poly-lysine-coated area, leaving a hydrophilic surface in the center (∼10 mm diameter). A piece of filter paper cut into a donut shape of 32 mm diameter with an 18 mm diameter hole was placed onto the bottom of the culture dish and made wet with water. Gravid hermaphrodite animals were dissected in 1 µl of dissection buffer (90 mM sucrose, 50 mM EGTA, 5 mM MgCl_2_, 50 mM potassium acetate, 50 mM PIPES-NaOH pH 7) dropped at the center of the poly-L-lysine coated area of the cover glass, immediately followed by addition of 100 µl of an isotonic medium based on Leibovitz’s L-15 medium (Gibco 21083-027) supplemented with 10% (v/v) fetal bovine serum, 35 mM sucrose, 100 units/mL penicillin and 100 µg/mL streptomycin (Christensen et al., 2002; Edgar, 1995). The sucrose concentration was optimized for the viability and normal embryonic divisions upon eggshell permeabilization by *perm-1* RNAi (Carvalho et al., 2011). To remove unattached embryos and the corpses of the mothers, the medium was replaced with another 100 µl of the same medium. The culture dish was then covered with a 50 mm diameter cover glass and mounted on the stage of an Andor Revolution XD spinning disk confocal microscopy system based on a Nikon Eclipse Ti inverted microscope equipped with a Nikon CFI Apochromat Lambda S 60×/1.40 NA oil-immersion objective lens, a spinning-disk unit (Yokogawa CSU-X1) and an Andor iXon Ultra EM-CCD camera. Images were acquired using Andor IQ3 software. Fluorophores were excited by laser lines at wavelengths of 488 nm for GFP or 561 nm for mCherry. The temperature of the sample, which was monitored by a FLIR One thermal imaging camera (FLIR Systems, Inc.), was maintained at 21 °C (**Figs. 1** to **3**) or at 25 °C (**Fig. 4**) by circulating cold air within an environmental chamber with a side panel removed. Embryos were staged by differential interference contrast (DIC) observation and the progress of mitosis was monitored every 5 s by observing the state of chromosomes with histone-mCherry (**Fig. 3**) or as dark zones in NMY-2::GFP (**Fig. 4**). For fast recording of NMY-2::GFP, image acquisition was started immediately after anaphase onset (**Fig. 3** and **4**) or 70 s later (**Fig. 3**) to capture a set of 5 z-slice images (100 ms exposure, 133 nm/pixel) with 0.5 µm z-steps every 0.83 s (**Fig. 1**), 0.72 s (**Fig. 2** and **3**) or 1.25 s (**Fig. 4**).

For DIC microscopy in **Supplementary Fig. 4,** the embryos were mounted between a cover glass and a 2% (w/v) agarose pad in 0.7× egg salt on an FCS2 cooling device (Bioptechs) set at 25 °C, and then filmed with an Olympus BX-51 upright microscope equipped with a UPlanSApo 100×/1.4 NA objective, DIC optics, and a CoolSNAP HQ2 CCD camera (Photometrics) controlled by MicroManager (https://www.micro-manager.org/) (Edelstein et al., 2014).

### Image analysis

The microscope images were processed and analyzed by custom scripts written with the macro language of Fiji/ImageJ (https://fiji.sc) (Schindelin et al., 2012) and R language (https://www.r-project.org/). The 4D images of NMY-2::GFP were deconvolved time frame by time frame with the DeconvolutionLab2 plugin (http://bigwww.epfl.ch/deconvolution/) (Sage et al., 2017) using the Richardson-Lucy algorithm with total-variation regularization and a point spread function calculated by the PSF generator plugin (Kirshner et al., 2013) with the Born and Wolf model. The coordinates of the cell periphery were determined frame by frame in the bleach-corrected, average z-projections of non-deconvolved images using the Trainable Weka Segmentation plugin (https://imagej.net/Trainable_Weka_Segmentation) (Arganda-Carreras et al., 2017). The positions of the spindle poles were determined by human visual detection of the weak NMY-2 localization to the spindle and the spindle poles in anonymized movies.

The velocity of the cytoplasmic NMY-2::GFP particles (**Fig. 1c**) was determined by making kymographs of all the trajectories in four anaphase embryos and measuring their gradients. For automated scoring in **Fig. 2d**, cytoplasmic NMY-2::GFP particles were detected in each time frame of the deconvolved and average z-projected movies by finding maxima within the boundary of each embryo using MaximumFinder in Fiji/ImageJ. The particles detected in two consecutive time frames less than 7 pixels apart, which correspond to movement at 1.295 µm/s, were stitched into a trajectory by using a custom R script, and overlaid on the original movies to be checked by visual inspection (not shown). Each trajectory was analyzed as a 2-dimensional biased Brownian motion by calculating mean square displacement to determine its velocity and diffusion coefficient by quadratic fitting. The trajectories that appeared in four or more time frames (2.88 s) and moved longer than 3 pixels (0.4 µm) at a velocity faster than 0.3 pixels/frame (55.4 nm/s) were scored for calculating the sum of the intensities of the moving particles at each time point.

For analysis of the temporal change of the cortical density and flow of NMY-2::GFP, first, the region 40 pixels (5.33 µm) inside and outside of the edge of the cell in the average z-projected image was straightened so that the periphery of the embryo that was traced counter-clockwise starting from the anterior tip was placed from left to right and thus the posterior tip ended at the center (**Supplementary Fig. 2**). The intensity of the cortical NMY-2::GFP signal, which appeared now as a horizontal line in the middle of the straightened “edge” image, was normalized so that the local background (outside of the cell) is 0 and the local cytoplasmic level is 1. The kymograph of the normalized density at the cell cortex was then generated by reslicing the stack of the time series of the straightened edge images with horizontal lines at intervals of one pixel and averaging the five consecutive best focal planes, which corresponds to the peak of the NMY-2::GFP signal at the cell periphery (cortex) of 0.67 µm width. After averaging across multiple embryos, the kymograph was folded back at the center (=the posterior tip) so that the anterior and posterior tips are placed on the left and right edges, respectively. For **Fig. 3**, the average kymograph from the dataset obtained from 0 to 140 s p.a.o. and that from 70 to 215 s p.a.o. were merged by linearly changing the blending ratio for the overlapping period (70 s to 140 s p.a.o.) after correction for photobleaching, which is more profound at the cortex than in the cytoplasm due to slower exchange with unbleached molecules.

One-dimensional particle image velocimetry (1D-PIV) was performed by a custom R script that compares the one-dimensional distribution pattern of the cortical NMY-2 signal along the cell periphery in a time frame with that of the next time frame. The spatial resolution of the kymograph of the NMY-2 signal was increased 5-fold (**Fig. 3**) or 10-fold (**Fig. 4**) by interpolation. The local velocity was determined as the spatial shift needed to maximize the cross-correlation between the windows of the size of 256 pixels (6.83 µm, **Fig. 3**) or 384 pixels (5.11 µm, **Fig. 4**) from the two consecutive time points. This was scanned along the cell periphery and repeated through the temporal dimension to make a kymograph of the flow of NMY-2 and the spatial resolution was set back to the original one by averaging. After averaging across multiple embryos, the kymograph of the flow was folded back so that the anterior and posterior tips are placed on the left and right edges, respectively. Positive (anterior to posterior) and negative (posterior to anterior) flow velocities were presented by pseudocoloring with green and magenta, respectively.

## Acknowledgements

We thank Behrooz Esmaeili for help in *C. elegans* maintenance and Rob Cross for useful comments on the text. This work was supported by the Wellcome Trust Senior Investigator Award (WT101885MA) to M.K.B., Cancer Research UK program grant (C19769/A11985) to M.M., Wellcome-Warwick Quantitative Biomedicine Programme (WQBP, Institutional Strategic Support Fund: 105627/Z/14/Z) Seed Funding to M.M. and WQBP Interdisciplinary Summer School scholarship to. A.W.

## Author contributions

B.C., M.K.B. and M.M. initiated the project. B.C., A.W. and M.M. generated the new *C. elegans* strains. B.C. and M.H. performed live imaging. M.M. performed image analysis. B.C., M.K.B. and M.M. prepared the manuscript. M.K.B. and M.M. supervised the project.

---

**Supplementary Fig. 1.**
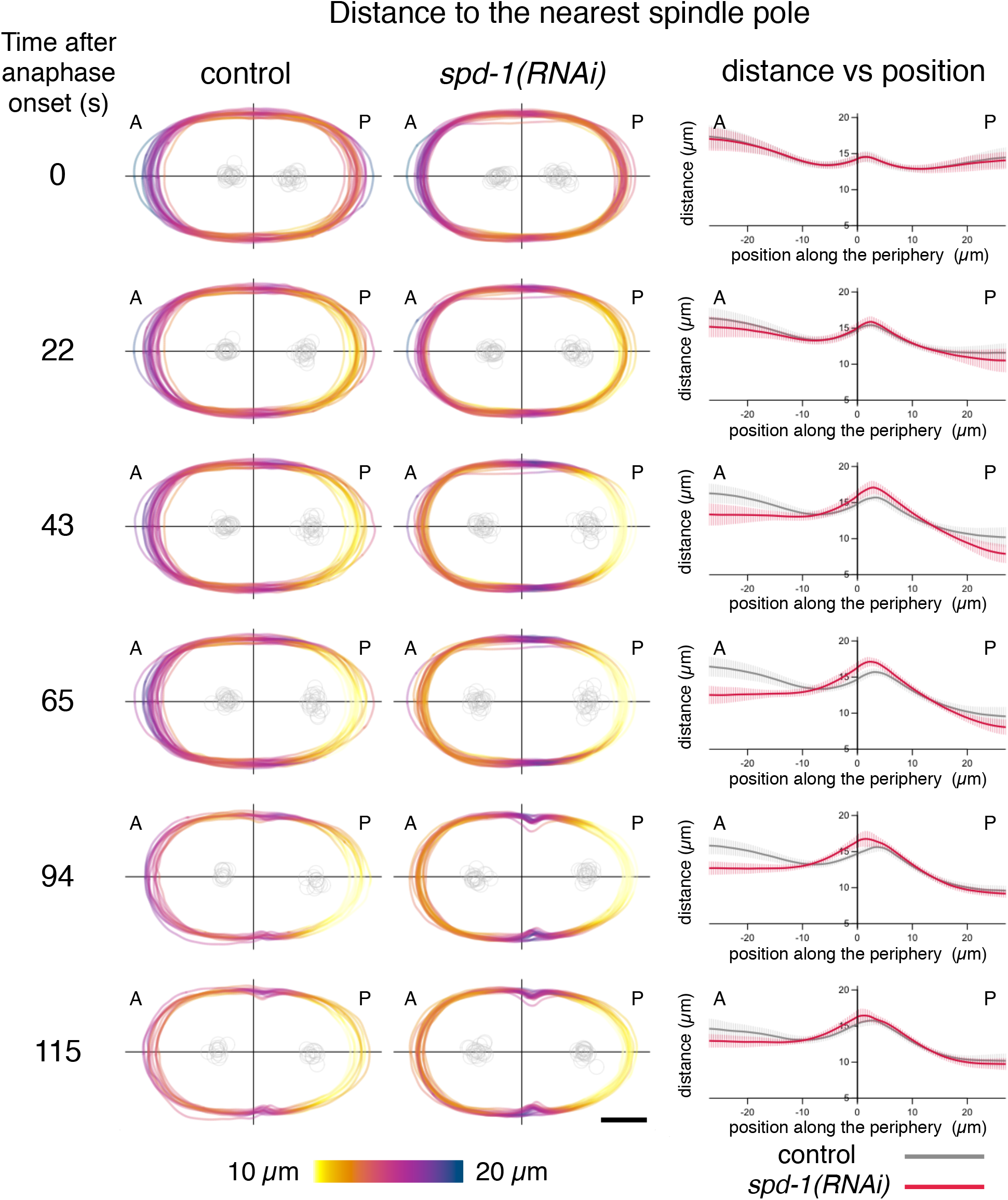
Distance from the cell cortex to the nearest spindle pole in *control(RNAi)* or *spd-1(RNAi)* embryos presented by color-coding and by plotting the distance vs position (0, 43 and 94 s are shown in Fig. 3a).

**Supplementary Fig. 2.**
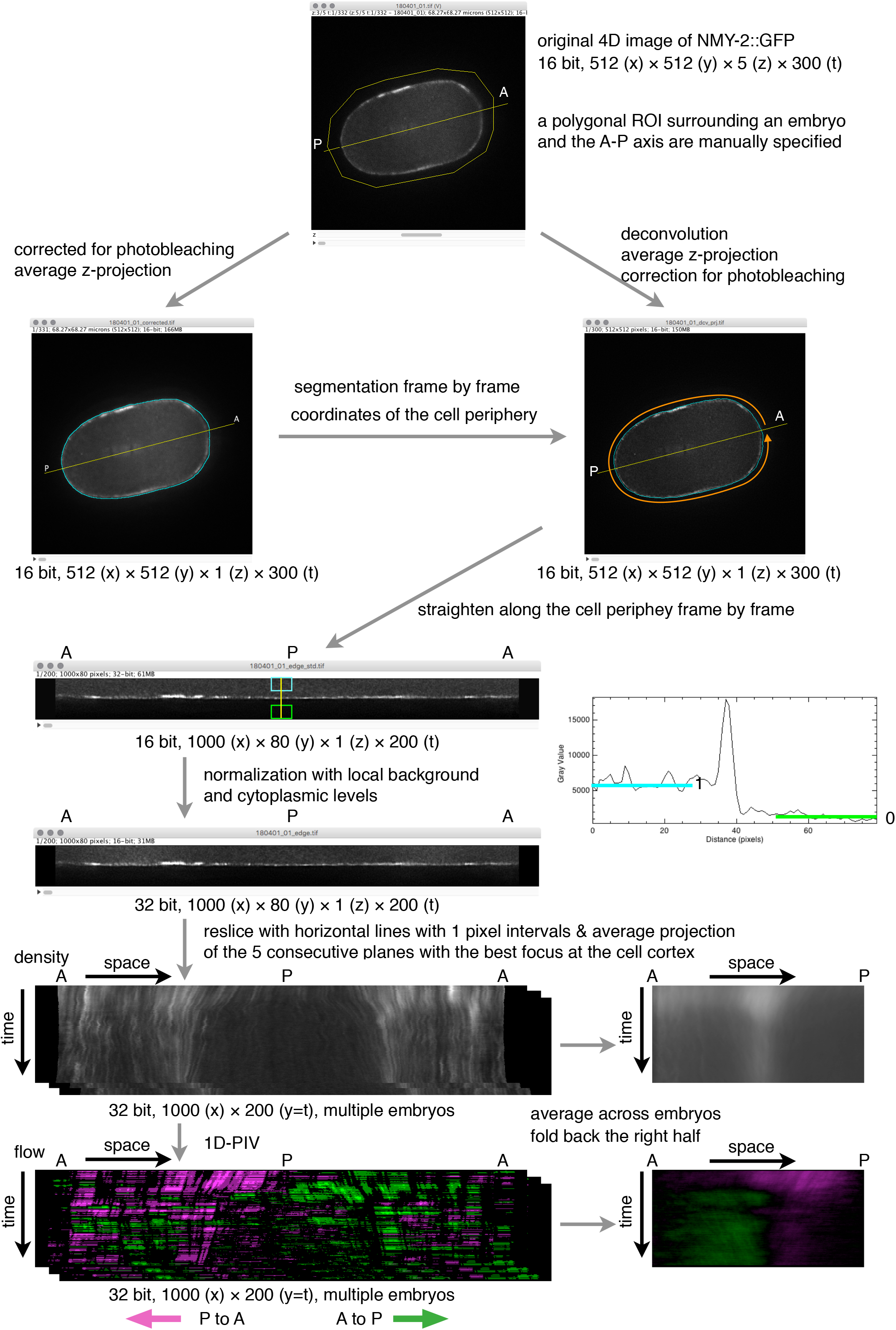
Schematic of the workflow of analysis of 4D image data of the *C. elegans* embryos expressing NMY-2::GFP. For more details, refer to Materials to Methods.

**Supplementary Fig. 3.**
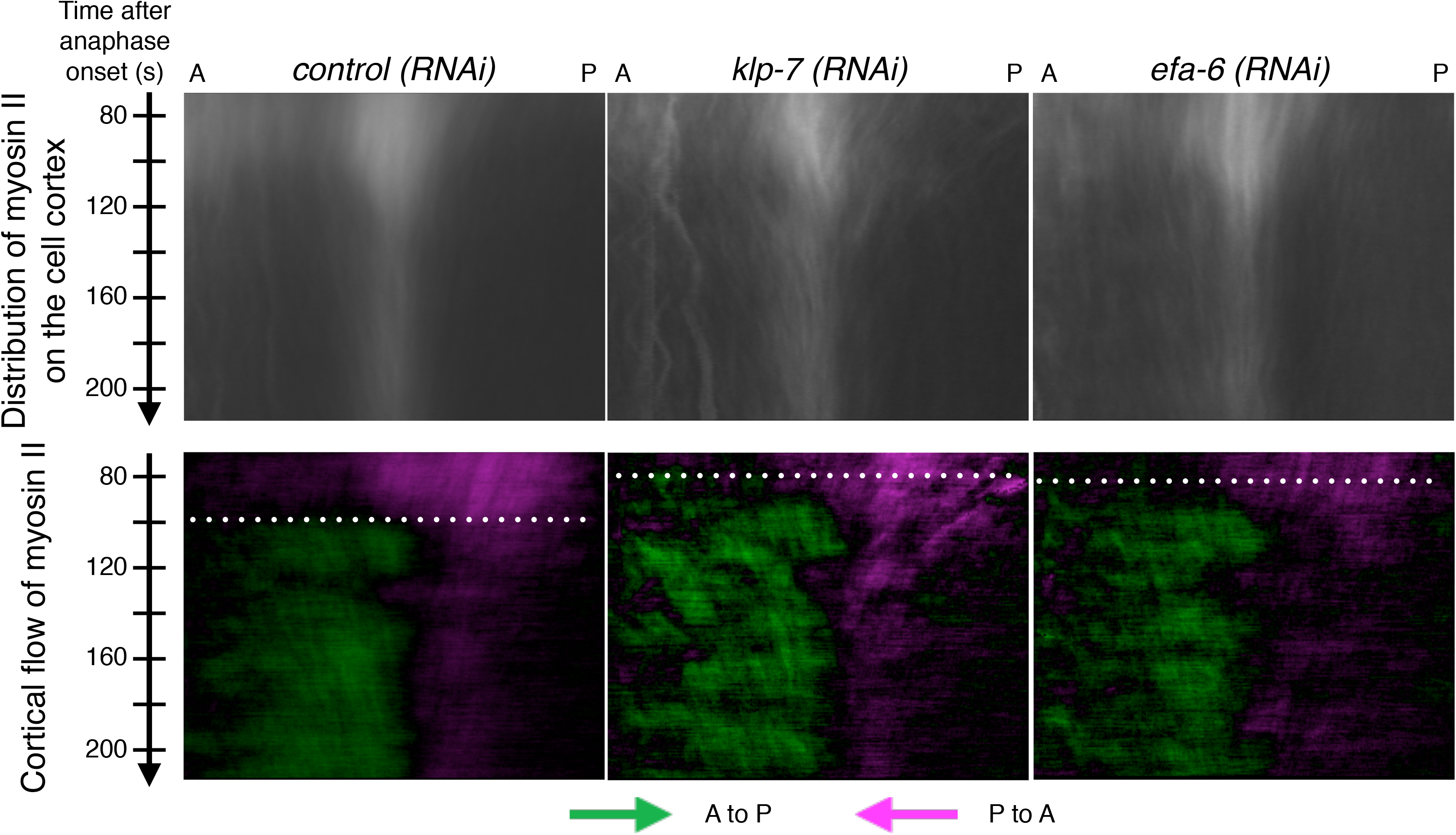
Distribution and flow of NMY-2::GFP in the cell cortex of the embryos depleted of KLP-7 (n=10) and EFA-6 (n=10). White dashed lines indicate the timings of the transition to the bidirectional mode of the cortical flow.

**Supplementary Fig. 4.**
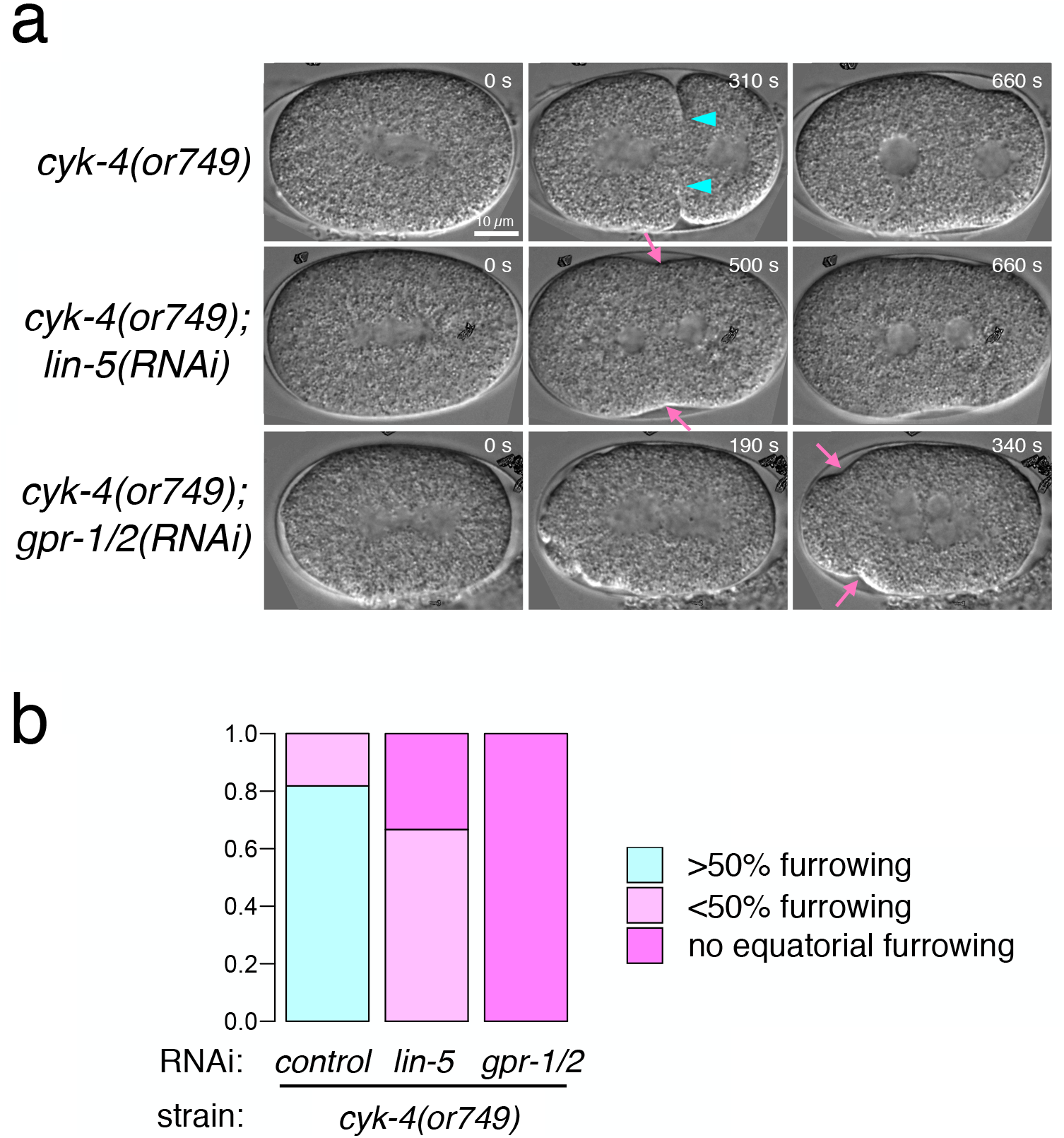
Synthetic effect of *cyk-4(or749)* and *lin-5(RNAi)* on the cleavage furrow formation. a. Stills from time-lapse recordings of differential interference contrast microscope images. 80% of the *cyk-4(or749)* embryos formed a cleavage furrow that showed 50% or deeper ingression (cyan arrowheads) but later regressed. Depletion of LIN-5 or GPR-1/2 aggravated this phenotype, allowing only shallow ingression (magenta arrows).

**Supplementary Table 1.** List of *C. elegans* strains used in this study

| Strain | Genotype | Reference |
| --- | --- | --- |
| LP162 | *nmy-2(cp13[nmy-2::gfp + LoxP]) I* | Dickson et al., 2013 |
| OD56 | *unc-119(ed3) III; ltIs37[(pAA64)*<br>*pie-1p::mCherry::his-58 + unc-119(+)] IV* | Essex et al., 2009 |
| QM160 | *nmy-2(cp13[nmy-2::gfp + LoxP]) I; ltIs37[(pAA64)*<br>*pie-1p::mCherry::his-58 + unc-119(+)] IV* | This study |
| EU1404 | *cyk-4(or749) III* | Canman et al., 2008 |
| QM196 | *nmy-2(cp13[nmy-2::gfp + LoxP]) I; cyk-4(or749) III* | This study |

**Supplementary Video 1**

NMY-2::GFP in the midplane of a *C. elegans* one-cell stage embryo during metaphase.

**Supplementary Video 2**

NMY-2::GFP in the midplane of a *C. elegans* one-cell stage embryo during anaphase.

**Supplementary Video 3**

NMY-2::GFP in the midplane of a one-cell stage *dhc-1(RNAi)* embryo in anaphase.

**Supplementary Video 4**

NMY-2::GFP in the midplane of a one-cell stage *lin-5(RNAi)* embryo in anaphase.

